# BCKDK Inhibitors Directly Antagonize Angiotensin II Type 1 Receptor and Induce Vasodilation Independently of BCAA Catabolism or BCKDK

**DOI:** 10.64898/2026.10.06.757051

**Authors:** Ryo Kawakami, Yann Deleye, Esra Akkaya, Uyen Pham, Jae Woo Jung, Ling Lai, Anand Chundi, Srikrishna Darbha, Danielle Murashige, Ryan B. Gaspar, Wentao He, Guo-Fang Zhang, Alem W. Kahsai, Sudarshan Rajagopal, Phillip J. White, Zoltan Arany, Robert W. McGarrah

## Abstract

Branched-chain amino acids (BCAA) have been implicated in diabetes and heart failure, leading to ongoing clinical trials with inhibitors of BCKDK, a key regulator of BCAA catabolism. We have previously shown that these inhibitors promote vasodilation and lower blood pressure, well-known cardioprotective effects, but how this occurs remained unclear. We show here, using explanted vessels, in vivo hemodynamics and telemetry, and genetically modified mice, that the vasodilatory effect of BCKDK inhibitors is direct, rapid, independent of nitric oxide, independent of BCAA catabolism, and most strikingly, independent of BCKDK itself. Instead, we demonstrate, using structural and biochemical studies, that BCKDK inhibitors directly antagonize the angiotensin II type I receptor (AT1R) and consequent angiotensin-mediated vasoconstriction. These data demonstrate a strong off-target effect of BCKDK inhibitors, independent of BCAA or BCKDK, which may explain their cardioprotective effects but also questions if this protection would be additive to existing AT1R blockers.

## Introduction

Branched-chain amino acids (BCAA: leucine, valine, and isoleucine) are essential amino acids that comprise up to 25% of protein in mammals. Elevations of plasma BCAA are a hallmark of heart failure (HF) and metabolic disorders such as obesity and diabetes. Downregulation of the enzymes involved in cardiac BCAA metabolism, and the accumulation of BCAA and their cognate α-keto acids (BCKA) in heart and circulation are now well-described features of both murine and human HF^1–3^. BCAA levels are in large part controlled by the catabolism of BCAA, the rate-limiting step of which is the dehydrogenation and decarboxylation of BCKA by the large BCKA dehydrogenase complex (BCKDH). BCKDH is inhibited by phosphorylation on specific residues mediated by BCKDH kinase (BCKDK). Inhibitors of BCKDK thus boost BCAA catabolism by the BCKDH complex and lower levels of BCAA and BCKA (Figure 1A). BT2, a widely used BCKDK inhibitor^4^, prevents maladaptive cardiac remodeling in numerous rodent HF models^5–9^, which has underscored the thesis that aberrant BCAA catabolism contributes to HF. Structural analogs of BT2 have been developed, one of which, PF-07328948 (PF’8948), is currently in a Phase II trial (NCT06991257) for the treatment of HF with preserved ejection fraction (HFpEF)^10,11^. The mechanism by which BT2 and PF’8948 are cardioprotective, however, remains unclear. Cardiac- or skeletal muscle-specific deletion of *Bckdk* in mice failed to provide cardioprotection^9^. We recently reported that BT2 induces vasodilation and lowers blood pressure (BP) in mice, suggesting that BCKDK inhibitors may instead confer cardioprotection via vasodilation^9^. However, how these agents promote vasodilation is unknown. We address this question in depth here.

**Figure 1.**
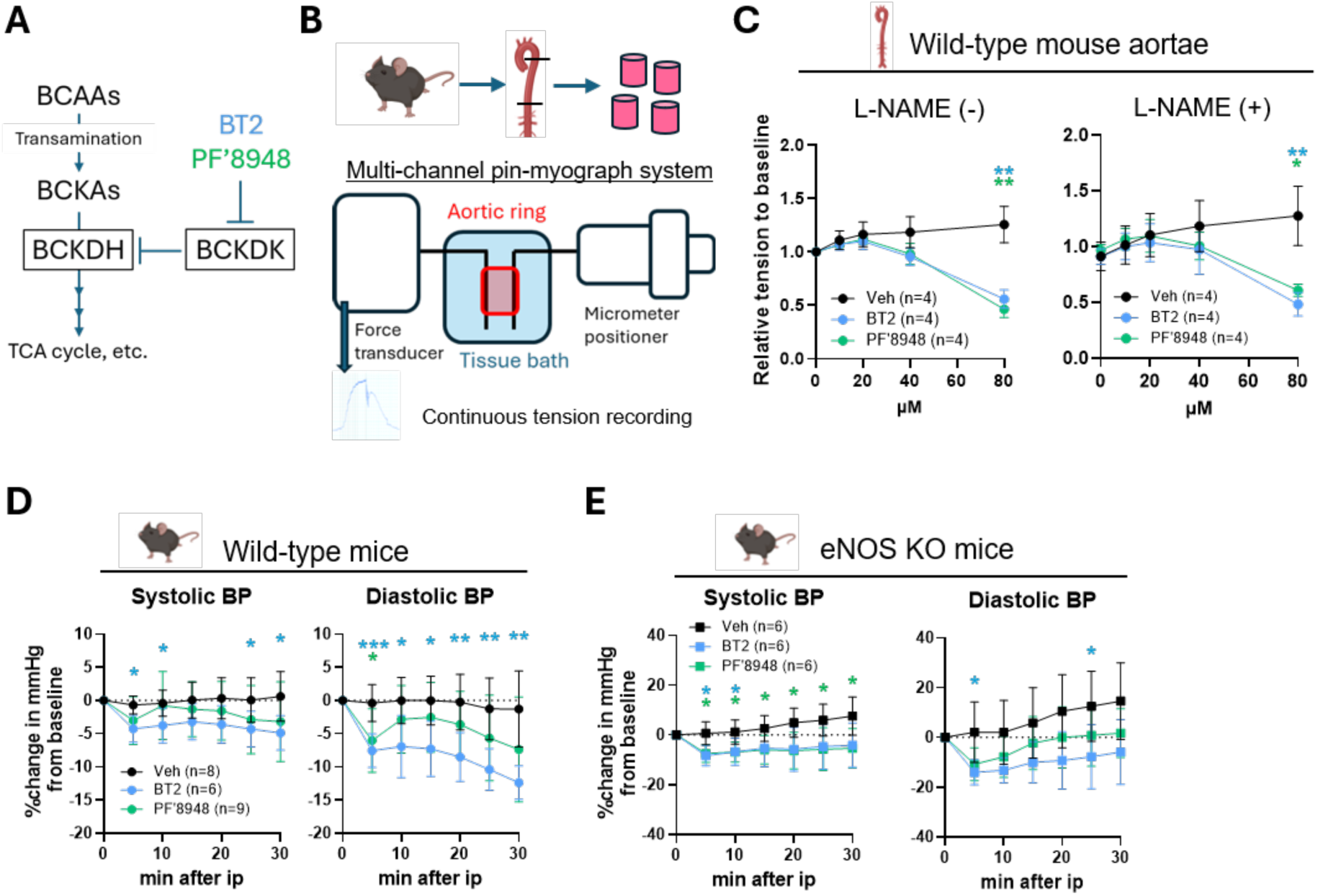
Direct, rapid, and NO-independent vasodilation by BCKDK inhibitors. **A)** A simplified diagram of the BCAA catabolic pathway. **B)** A multi-channel pin-myograph system (DMT 820MO) records isometric tension of vascular rings, isolated from mouse thoracic descending aorta, prestretched to 0.5-gram tension. **C)** Dose-response of phenylephrine (PE, 100 nM)-induced contraction to BT2 or PF-07328948 (PF’8948), with or without 30 min pretreatment with 100 µM L-NAME. **D)** A pressure-volume catheter (MillarPVR-1045) was placed into the aorta via the right carotid artery in isoflurane-anesthetized mice. BT2 (40 mg/kg) or PF’8948 (60 mg/kg) was injected intraperitoneally, and blood pressure (BP) was monitored for 30 min. **E)** The same BP monitoring was performed using whole-body eNOS KO mice. Data shown as Mean±SD. Repeated-measures 2-way ANOVA, followed by Dunnett’s multiple comparisons test, was performed between BT2/PF’8948 and vehicle (*p<0.05, **p<0.01, and ***p<0.001).

## Results

### Direct, rapid, and NO-independent vasodilation by BCKDK inhibitors

To begin elucidating the vascular effects of BCKDK inhibitors, we assessed vascular tone in isolated mouse aortae treated with either BT2 or PF’8948, using a multi-channel pin-myograph system (Figure 1B). Both compounds significantly reduced phenylephrine (PE)-induced contraction in a dose-dependent manner, and at doses typically used in other studies^4,9,12^ (Figure 1C). These agents thus (1) directly affect vascular tone, and (2) do so independent of systemic neurohormonal milieu. Of note, the vasodilatory responses occurred within a minute (Figure S1A), suggesting rapid post-transcriptional and post-translational effects.

Vasodilation is largely regulated by endothelial release of nitric oxide (NO) by endothelial NO synthase (eNOS). Surprisingly, however, vasodilation triggered by BT2 or PF’8948 was not blocked by suppressing eNOS with L-NAME (Figure 1C, right panel). To examine these effects *in vivo*, we administered BT2 or PF’8948 to wild-type and eNOS^-/-^ mice and monitored BP using an intravascular pressure catheter. Both compounds within 5 minutes of intraperitoneal injection lowered both systolic and diastolic BP, to various degrees, consistent with rapid vasodilatory effects (Figure 1D). Moreover, systolic and diastolic BP reductions were similarly seen in eNOS^-/-^ mice (Figure 1E). We conclude from these studies that these BCKDK inhibitors exert rapid and direct vasodilatory effects *in vivo* and *ex vivo* and do so independently of endothelial NO production.

### BCKDK inhibitors promote vasodilation independently of smooth muscle BCAA catabolism or of BCKDK itself

These data suggested a possible role of BCAA catabolism in smooth muscle (SM), rather than endothelium, in regulating vascular tone. To test this notion, we generated SM-specific inducible BCKDK knockout (SM-BCKDK KO) mice by providing *Bckdk*^lox/lox^:: *Myh11*-Cre ^ERT2^ mice with tamoxifen chow (Figure 2A). *Bckdk* deletion was efficient, leading to dephosphorylation and thus disinhibition of BCKDH (Figure 2A, right panels) and consequently increased ^13^C incorporation from U-^13^C-leucine to the TCA cycle in explanted aortae (Figure 2B), i.e. boosted BCAA catabolism. However, vascular reactivity of explanted aortae from SM-BCKDK KO mice was unaltered, compared to aortae from control mice (Figure 2C). Moreover, BP was no different in SM-BCKDK KO mice versus controls, even with several subsequent hemodynamic and vascular challenges such as high fat diet, high salt diet, and supplementation with L-NAME (Figure 2D). These findings indicate that enhanced BCAA catabolism in SM is not responsible for the vasodilatory effects of BT2 and PF’8948.

**Figure 2.**
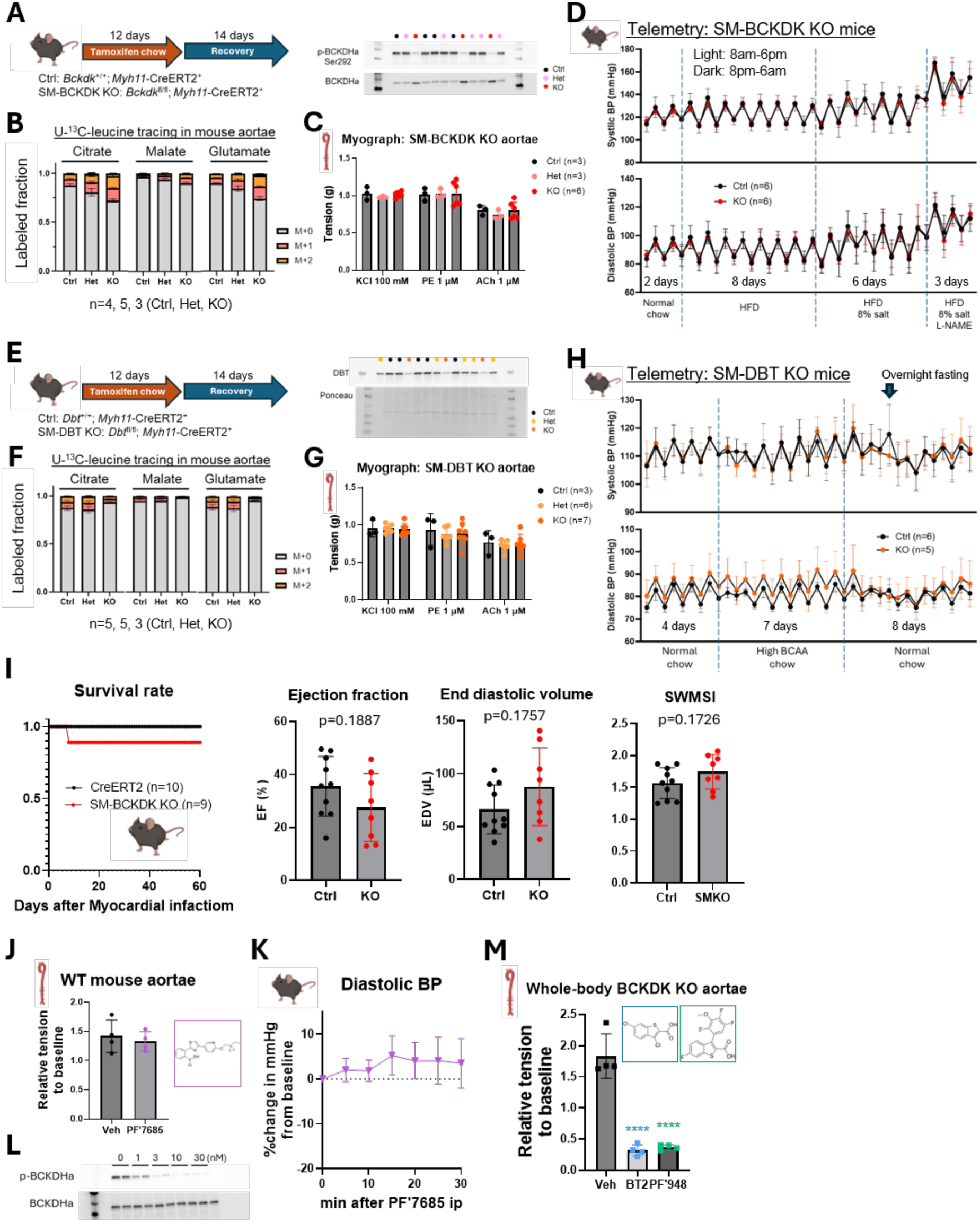
BCKDK inhibitors promote vasodilation independently of smooth muscle BCAA catabolism or of BCKDK itself. **A)** The scheme of generation of Smooth muscle (SM)-specific BCKDK KO mice, and Western blot of mouse aorta lysates to confirm KO. **B)** Mouse thoracic descending aortae were incubated in DMEM media containing U-^13^C-leucine for 3 hours. Metabolites extracted with methanol, acetonitrile, and water (2:2:1) were analyzed with liquid chromatography-mass spectrometry, and ^13^C-labeled fraction was calculated. **C)** Myograph with potassium chloride (KCl), PE, and acetylcholine (ACh). **D)** Telemetry (DSI HD-X11) monitored BP (high fat diet: Research Diets D12492; L-NAME 0.5 g/L in drinking water). Each dot represents average BP during the light or dark cycle. **E-H)** The same set of experiments were performed using SM-DBT KO mice. High BCAA chow contained 2x BCAA. **I)** SM-BCKDK KO mice subjected to left anterior descending artery ligation and echocardiography (VisualSonics Vevo 2100, at 8 weeks) in a blinded manner. Segmental Wall Motion Score Index (SWMSI) was determined to estimate the infarct size. Unpaired t-test was performed to compare Ctrl and KO mice. **J)** Myograph with WT mouse aortae and 10 µM PF-07247685 (PF’7685). **K)** BP monitoring in WT mice after 10 mg/kg PF’7685 injection. **L)** Western blot of mouse aorta lysates treated with PF’7685. **M)** Myograph with aortic rings isolated from whole-body BCKDK KO mice with BT2 or PF’8948 (80 µM each). Mice on a C57BL/6J background (male, 8-12 weeks) were used throughout. Data shown as Mean±SD. One-way ANOVA, followed by Dunnett’s multiple comparisons, was performed between BT2/PF’8948 and vehicle (****p<0.0001).

To further examine the physiological role of BCAA catabolism in SM, we also generated SM-specific loss of DBT, a critical subunit of the BCKDH complex (*Dbt*^lox/lox^:: *Myh11*-Cre ^ERT2^, hereafter SM-DBT KO mice, Figure 2E). Loss of SM DBT completely blocked ^13^C incorporation from U-^13^C-leucine to the TCA cycle in explanted aortae, demonstrating loss of BCAA oxidation (Figure 2F). However, vascular reactivity (Figure 2G) and BP (Figure 2H) were no different between SM-DBT KO and control mice, whether receiving regular or high BCAA-containing chow, again indicating a minor role, if any, of BCAA catabolism in regulating vascular tone.

These data raised the possibility that cardioprotection by BCKDK inhibition is mediated by SM BCKDK inhibition but not vasodilation. To investigate this possibility, we subjected SM-BCKDK KO mice to myocardial infarction and evaluated subsequent cardiac maladaptation. However, we saw no significant difference in survival rates, cardiac function, or estimated infarct size between the SM-BCKDK KO mice and controls (Figure 2I and Figure S2). We conclude that inhibition of SM BCKDK is not cardioprotective, and thus unlikely to mediate the cardioprotective effects of BT2 and PF’8948.

These unexpected data raised the alternate possibility that the vasodilatory effects of BCKDK inhibitors are mediated by a target other than BCKDK itself. To address this possibility, we first tested a different, and more potent, BCKDK inhibitor (PF-07247685 [PF’7685]), which importantly is structurally distinct from BT2 and PF’8948^10^. PF’7685 neither caused vasodilation *ex vivo* (Figure 2J) nor affected BP in mice *in vivo* (Figure 2K), despite almost complete loss of BCKDHA phosphorylation (Figure 2L), demonstrating that BCKDK inhibition is insufficient to cause vasodilation. We next isolated aortae from whole-body *Bckdk*^-/-^ mice and tested the effects of BT2 and PF’8948. Both compounds showed clear vasodilation of aortae from *Bckdk*^-/-^ mice (Figure 2M). These findings thus unequivocally and surprisingly demonstrate that BCKDK inhibition is neither necessary nor sufficient to cause vasodilation, and that the vasodilatory effects of BT2 and PF’8948 occur independently of BCKDK.

### BCKDK inhibitors antagonize angiotensin II-mediated vasoconstriction and bind to the orthosteric site of AT1R

We therefore next sought to investigate the mechanism underlying the observed vasodilation. Recent studies have shown that angiotensin receptor blockers (ARBs), medications used in the treatment of hypertension and HF, bind to the allosteric regulatory pocket of BCKDK, and inhibit its activity in a manner similar to BT2^13^. We therefore postulated that, reciprocally, BT2 might bind to the target of ARBs, the angiotensin II type 1 receptor (AT1R), and inhibit angiotensin signaling to lower BP and provide cardioprotection independent of its effects on BCAA metabolism. Consistent with this notion, both BT2 and PF’8948 significantly reduced angiotensin II-induced contraction of mouse abdominal aortae, in the presence of L-NAME, while the structurally distinct BCKDK inhibitor, PF’7685, did not (Figure 3A).

**Figure 3.**
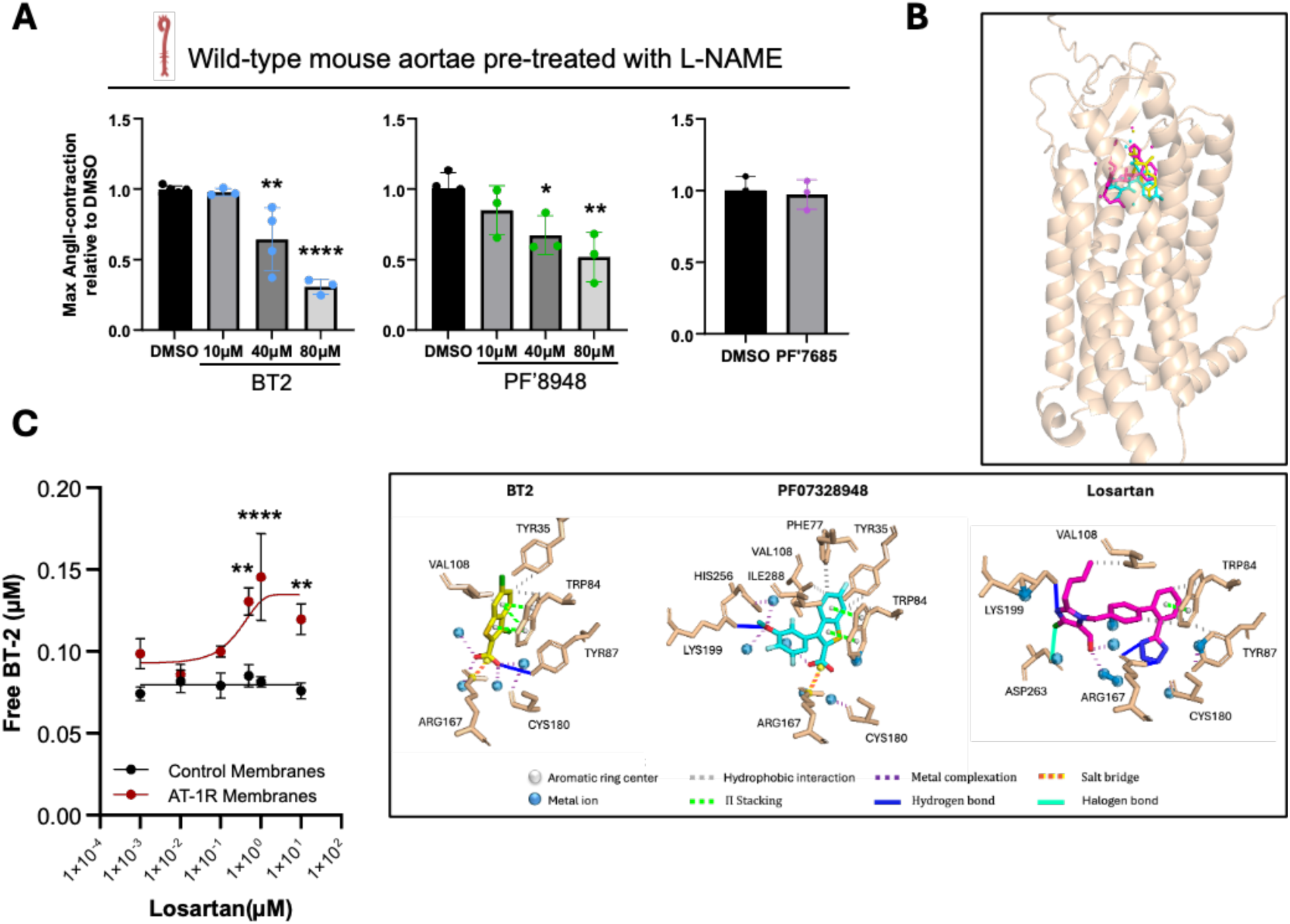
BCKDK inhibitors antagonize angiotensin II-mediated vasoconstriction and bind to the orthosteric site of AT1R. **A)** A multi-channel pin-myograph system (DMT 820MO) records isometric tension of vascular rings, isolated from mouse infra-renal aorta, prestretched to 0.4-gram tension. The rings were pre-treated with 100 µM L-NAME for 15 min and with BT2 (10-80 µM), PF’8948 (10-80 µM), or PF’7685 (10 µM) for 5 min, followed by 100 nM angiotensin II-induced contraction. One-way ANOVA, followed by Dunnett’s multiple comparisons, was performed between BT2/PF’8948/PF’7685 and vehicle (*p<0.05, **p<0.01, and ****p<0.0001). **B)** Top panel: Cartoon representation of a lateral view of the Alphafold-predicted structure of human AT1R, with BT2 (yellow), PF’8948 (cyan) and Losartan (magenta) simulated docking in the orthosteric site. Bottom panel: PLIP interactions between BT2 (yellow), PF’8948 (cyan), Losartan (magenta) with the key interacting residues of AT1R orthosteric site. **C)** Displacement of BT2 from AT1R by increasing doses of losartan (**p<0.01, ****p<0.0001).

Molecular modeling of BT2 with AT1R in the active conformation revealed the potential for BT2 to bind to the orthosteric pocket of AT1R in a manner similar to ARBs (Figure 3B). BT2 establishes three critical interactions in the orthosteric site (BT2 binding affinity -7.636 kcal/mol) that are consistent with ARB activity (losartan binding affinity -10.98 kcal/mol). These include the formation of salt bridges between Lys199 of transmembrane domain 5 and the carboxylic group of BT2, hydrophobic interaction (pi–cation) of BT2 with Arg167 on the ligand-binding pocket side of extracellular loop 2, and hydrophobic interaction (pi–pi) of BT2 with Trp84 of transmembrane domain 2 (Figure 3B). Importantly, most ARBs establish contacts with Arg167 and Trp84, while telmisartan and candesartan additionally contact Lys199. Similarly, critical interactions were also predicted between BT2 and AT1R when modeled in the inactive conformation (Supplemental Figure 3A). Furthermore, modeling of the interaction of PF’8948, which is structurally similar to BT2, with AT1R predicted its binding to the AT1R orthosteric pocket in a manner similar to BT2 and ARBs (PF’8948 binding affinity -8.818 kcal/mol) (Figure 3B), while PF’7685, which is structurally distinct from BT2 and PF’8948, was not predicted to bind.

To experimentally test interaction of BT2 with the ARB binding domain in the orthosteric pocket of AT1R, we incubated membranes from Expi293 cells overexpressing recombinant AT1R (Supplemental Figure 3B) with BT2 and measured the displacement of BT2 from the membrane upon exposure to increasing doses of losartan (Figure 3C). Exposure of the BT2-bound membranes to increasing doses of losartan led to increased concentrations of displaced BT2 in the media, consistent with direct competition of ARB with BT2 binding to the orthosteric pocket of AT1R. This effect was not observed in control membranes lacking AT1R (Figure 3C).

### BT2 inhibits AT1R signaling

We next examined whether BT2 modulates angiotensin II (AngII)-dependent AT1R signaling in HEK293T cells, using well-established assays previously described^14^. BT2 pretreatment reduced AngII-induced AT1R internalization in a concentration-dependent manner, as measured by BRET between AT1R-Rluc8 and the endosomal marker 2×FYVE-mVenus (Figure 4A). Consistent with impaired receptor trafficking, BT2 also decreased AngII-stimulated β-arrestin 1 recruitment to AT1R, assessed using the NanoBiT complementation assay (Figure 4B). Because AT1R signaling also engages G-protein-dependent calcium mobilization, we evaluated intracellular calcium responses following AngII stimulation. BT2 pretreatment similarly attenuated the calcium influx in a concentration-dependent manner (Figure 4C). Together, these findings show that BT2 suppresses multiple downstream consequences of AT1R activation, including β-arrestin recruitment, receptor internalization, and calcium signaling.

**Figure 4.**
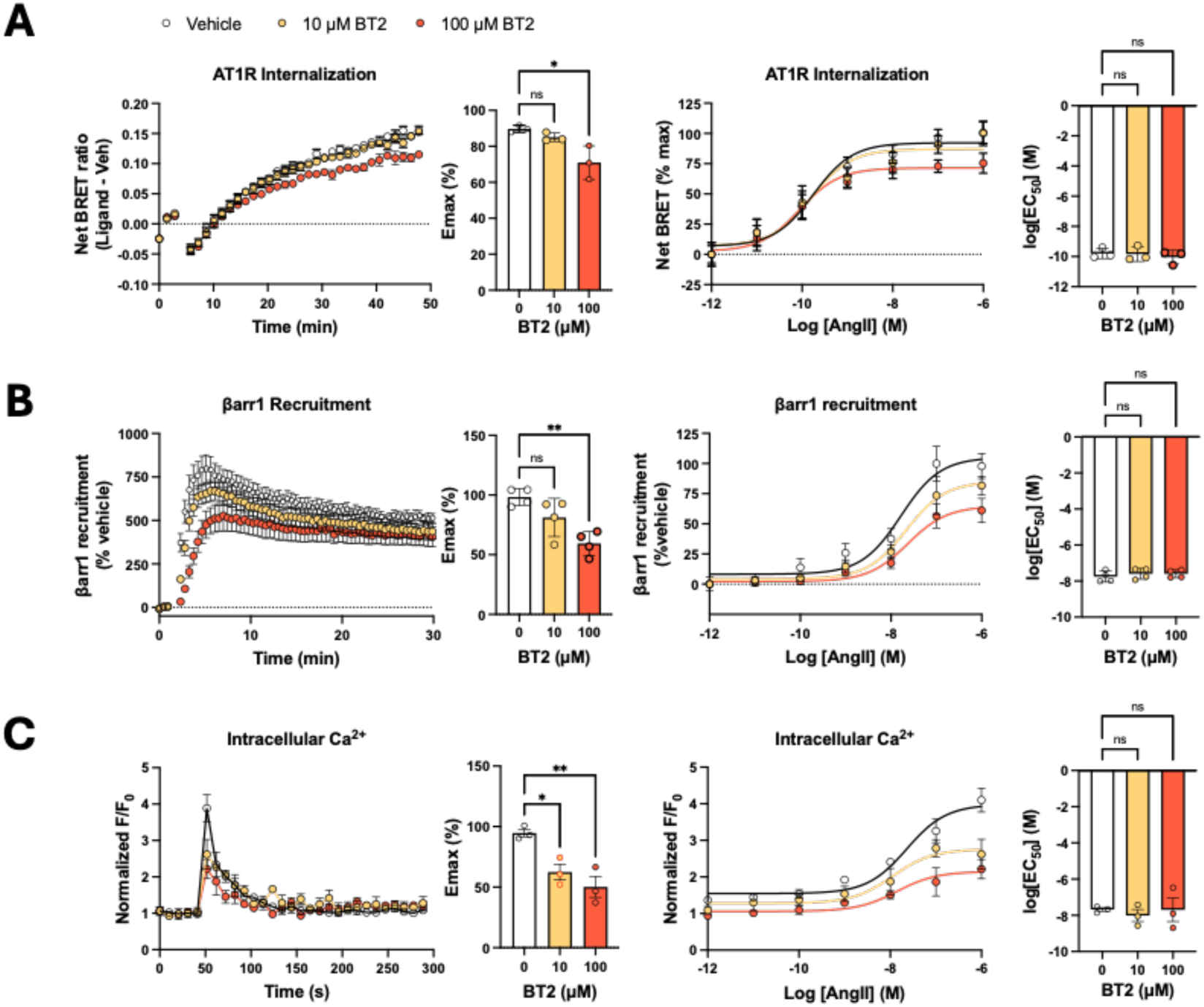
BT2 decreases AngII-mediated AT1R internalization, β-arrestin 1 recruitment, and intracellular calcium influx. HEK293T cells expressing AT1R were pretreated with BT2 or vehicle before stimulation with dose response of AngII. **A**) AT1R internalization, measured by BRET between AT1R-Rluc8 and the endosomal marker 2×FYVE-mVenus (n = 3 independent experiments). **B**) β-arrestin 1 recruitment to AT1R, measured using the NanoBiT assay (n = 4 independent experiments). **C**) AT1R-mediated intracellular calcium mobilization measured with Cal-520 AM (n = 3 independent experiments). Left panels show kinetic responses following AngII stimulation; right panels show AngII concentration-response curves. BT2 reduced the maximal AngII-evoked response in each assay without an apparent change in AngII potency. Data represent mean ± SEM. Statistical comparisons were performed by one-way ANOVA with Dunnett’s post hoc test for BT2 versus vehicle-treated cells. *P < 0.05; **P < 0.01; ns, not significant.

To assess receptor selectivity, we tested BT2 in cells expressing the vasopressin V2 receptor (V2R). BT2 did not significantly alter maximal agonist-dependent V2R β-arrestin recruitment or internalization (Supplemental Figure 3C), indicating that its inhibitory effects are selective for AT1R under these experimental conditions.

## Discussion

Our findings identify an unexpected mechanism underlying the vascular effects of BT2 and PF’8948 that is independent of their intended target, BCKDK. Instead, the data support direct antagonism of AT1R by BT2, providing a mechanistic link between these compounds, vasodilation, and reduced blood pressure. Recent studies reported other off-target effects of BT2, including mitochondrial uncoupling^15^ and enhanced tryptophan catabolism^16^. However, neither likely explains the vasodilation and cardioprotection conferred by the BCKDK inhibitors. In contrast, our present findings provide a mechanistic explanation for both the vasodilatory properties of the compounds, via action in the vascular smooth muscle and intracellular calcium influx, and their cardioprotective properties, via afterload reduction and perhaps diminished angiotensin II signaling in other cells as well. It will be of future interest to test these compounds in Agtr1a^-/-^ mice or vessels, to test if AT1R inhibition is the exclusive mediator of the impact by these compounds on vasodilation and cardioprotection.

We show here that the vasodilatory effects by BT2 and PF’8948 are distinct from any changes that these compounds exert on BCAA catabolism or protein phosphorylation networks through BCKDK inhibition. This finding likely explains why BCKDK deletion in heart, skeletal muscle, liver, and smooth muscle, has all failed to confer cardioprotection in various models of HF^5–9^.

Indeed, many conclusions in the literature about the role of BCAA in various contexts have relied heavily on studies with BT2 and should be reevaluated in light of our current findings. For example, it is possible that BT2-mediated vasodilation, independent of any effect by BCAA, promotes microvascular flow and thus insulin sensitivity in skeletal muscle. Our results thus provide a cautionary tale for mechanistic studies overly reliant on small molecule inhibitors, which are almost always pleiotropic, and often in ways that are not known.

Our work also has important implications for ongoing and future clinical trials with this class of BCKDK inhibitors. It may be, for example, that clinical effects of these compounds do not differ substantially from those conferred by ARBs, now generic and widely used. Such trials should therefore consider concurrent use of ARBs to determine if BCKDK inhibition provides additive clinical benefits.

## Article Information

## Acknowledgement

The catheter implantation for BP monitoring and the myocardial infarction surgery, followed by echocardiography, were performed by the Rodent Cardiovascular Phenotyping Core (RRID: SCR_022419) at the University of Pennsylvania supported by the Penn Cardiovascular Institute and NIH S10OD016393. We thank the University of Pennsylvania Rodent Metabolic Phenotyping Core (RRID: SCR_022427) for the telemetry implantation surgery, supported in part by NIH grant S10-OD025098 and the Cox Institute. We thank Laura Wingler, PhD, for sharing the AT1R overexpression plasmid for AT1R binding assays.

## Methods

### Animal models

All animal procedures were approved by the Institutional Animal Care and Use Committee (IACUC) of the University of Pennsylvania and were performed in accordance with approved protocols. Mice were maintained on a 12-h light/12-h dark cycle under controlled temperature and humidity conditions, with ad libitum access to food and water. Whole-body *Bckck*-knockout, *Bckdk*^fl/fl^, and *Dbt*^fl/fl^ mice were generated as previously described^9,17^. Mice carrying the respective floxed alleles were then crossed with Myh11-CreERT2 mice (B6.FVB-Tg(Myh11-icre/ERT2)1Soff/J; The Jackson Laboratory, stock no. 019079). Smooth muscle-specific gene deletion was induced by feeding the mice tamoxifen-containing chow (Envigo, TD.130856) for 12 days^18^. Mice were subsequently returned to standard chow (Purina LabDiet 5010) for at least 14 days before experimentation to allow tamoxifen clearance. Genetically modified mouse lines were maintained on a C57BL/6J background. Tamoxifen-treated, Myh11-CreERT2-positive littermates that were wild-type at the targeted locus served as controls. Because the Myh11-CreERT2 transgene is integrated into Y chromosome, only male mice carry the transgene. For myograph experiments involving wild-type or eNOS-knockout mice (Figure 1, 2K, and 3A), mice were purchased from Jackson laboratory (C57BL6J: #000664 and B6.129P2-Nos3tm1Unc/J: #002684). Male mice and 8–12 weeks of age were used unless otherwise specified.

### Pin-myograph

Vasoreactivity was measured using a four-channel pin-myograph system (DMT 820MO; Danish Myo Technology, Denmark) according to standard procedures^19^. Mice were euthanized by isoflurane overdose, and the descending thoracic or infrarenal abdominal aortae were immediately excised. Surrounding adipose and connective tissues were removed under a dissecting microscope in ice-cold physiological salt solution (PSS; NaCl 130 mM, KCl 4.7 mM, KH2PO4 1.18 mM, MgSO4 7H2O 1.17 mM, NaHCO3 24.9 mM, Glucose 5.5 mM, EDTA 260 µM, CaCl2 1.6 mM). Thoracic and abdominal aortae were cut into rings 2.0 and 1.5 mm in length, respectively, and mounted on the pins in myograph chamber containing 5 mL of warmed PSS. The PSS was continuously aerated with 95% O_2_/5% CO_2_, and maintained at 37 °C. During a 45-min equilibration period, the rings were gradually stretched to a resting tension of 0.5 g for thoracic aortae or 0.4 g for abdominal aortae, with three changes of PSS. Contractile responsiveness was confirmed by exposure to 100 mM KCl for 5 min, and the peak KCl-induced contraction was used to normalize subsequent contractile responses. The chambers were then washed three times with 8mL of PSS per wash, and the rings were allowed to re-equilibrate for 30 min, with one additional change of PSS. BCKDK inhibitors were dissolved in DMSO, whereas other compounds were dissolved in water, to prepare 500× or 1,000× stock solutions. Depending on the stock concentration, 5 or 10 µL was added to each 5-mL myograph chamber to achieve the indicated final concentration. An equal amount of the corresponding vehicle was added to each control chamber. Phenylephrine or angiotensin II was then added to induce contraction, and peak contractile response were compared between treatment groups. Data were acquired and analyzed using LabChart 8 software (ADInstruments).

### Acute invasive blood pressure measurement

Hemodynamic measurements by carotid artery catheterization were performed by the Rodent Cardiovascular Phenotyping Core (RRID: SCR_022419) at the University of Pennsylvania. The mice were anesthetized with 2.5% inhaled isoflurane, intubated, and mechanically ventilated using a Harvard ventilator set to deliver a tidal volume of 200–400 µL. The right carotid artery was isolated and cannulated with a pressure-volume catheter (Millar PVR1045) for continuous arterial pressure monitoring. Systolic, diastolic, and mean arterial pressures were recorded and analyzed using a PowerLab 30 system (ADInstruments, Colorado Springs, CO, USA). BT2, PF-07328948, and PF-07247685 were formulated at concentrations of 10, 15, and 0.25 mg/mL, respectively, in a vehicle consisting of 5% DMSO, 10% Cremophor EL (MilliporeSigma, catalog no. 238470), and 85% 0.1 M sodium bicarbonate adjusted to pH 9.0 (v/v/v). Compounds were administered intraperitoneally at 4 µL/g body weight, corresponding to doses of 40, 60, and 10 mg/kg, respectively. Vehicle-control mice received an equivalent volume of vehicle.

### Western blot

Aortic tissue was homogenized in 1× RIPA buffer supplemented with phosphatase and protease inhibitors (PhosSTOP and Complete mini proteinase inhibitor cocktail, Roche). Approximately 5 mm of aorta was homogenized in a 2 mL plastic tube containing 100 µL of RIPA buffer and 1.5 mm ceramic beads using Bead Ruptor 12 Bead Mill Homogenizer (Omni; Speed 4 m/s, two 40-s cycles). The lysates were centrifuged at 16,000 g for 10 min and the supernatant was collected. The protein concentration was quantified using a BCA assay kit (Thermo Fisher Scientific). Samples containing 3.5 µg of protein in a total volume of 15 µL were prepared with 4x Laemmli buffer containing 2-mercaptoethanol, heated at 95C for 5 min, and separated by SDS-PAGE using using 4–20% Criterion TGX Precast Midi Protein Gels (26-well, 15-µL format; Bio-Rad, catalog no. 5671095). Proteins were transferred to a PVDF membrane, and the membrane was incubated with primary antibodies against phospho-BCKDHa (Ser293; Bethyl Laboratories, A304-672A, 1:1,000), BCKDHa (Bethyl Laboratories, A303-790A, 1:1,000), and DBT (Invitrogen, PA5-29727, 1:1,000) and an HRP-conjugated anti-rabbit IgG secondary antibody (Cell Signaling Technology, catalog no. 7074, 1:5,000). Pierce™ Protein-Free Blocking Buffer (Thermo Scientific, catalog no. 37571) was used for blocking membranes and diluting the antibodies. Signals were detected using Pierce SuperSignal Femto or Pico PLUS chemiluminescent substrate and a CCD-based imaging system.

### Stable isotope tracing

Mice were euthanized by isoflurane overdose, and the descending thoracic aortae were excised. The surrounding adipose and connective tissues were removed in ice-cold RPMI 1640 under a dissecting microscope. Aortic rings 4 mm in length were pre-incubated in warm RPMI 1640 in a CO2 incubator at 37 °C for 30 min. The medium was replaced with BCAA-free RPMI 1640 supplimented with 300 µM valine, 200 µM uniformly ^13^C-labeled leucine, and 100 µM isoleucine. After 3 h, the aortic rings were rapidly removed from the medium, briefly blotted to remove residual medium, and immediately transferred into 50 µL of extraction solvent (methanol, acetonitrile, and water [2:2:1]). The extracted metabolites were measured by liquid chromatography coupled to a Q Exactive mass spectrometer (ESI, negative ion mode, full scan MS). XBridge Premier BEH Amide (2.5 µm, 2.1 x 150 mm) was used under basic HILIC conditions. Mobile phases A consisted of 20 mM ammonium acetate and 20 mM ammonium hydroxide in water/acetonitrile (95:5, v/v), adjusted to pH 9.5, and mobile phase B consisted of 100% acetonitrile. A 5-µL aliquot of each sample was injected using an autosampler. The LC gradient was as follows: 0–2 min, 90% B; 3–5 min, 80% B; 6–7 min 75% B; 8–9 min, 70% B; 10–11 min, 50% B; 12–14 min, 40% B; 15–20 min, 90% B. Data were analyzed using El-MAVEN.

### Long-term blood pressure monitoring by radiotelemetry

Surgical implantation of HD-X11 radiotelemetry transmitters (Data Sciences International, St. Paul, MN, USA) was performed by the Rodent Metabolic Phenotyping Core (RRID: SCR_022427) at the University of Pennsylvania. The catheter was inserted through the right carotid artery and advanced into the aorta. Following implantation, mice were individually housed in a dedicated telemetry room with ad libitum access to food and water. Mice were allowed to recover for 1 week before data collection. Arterial pressure waveforms were continuously acquired using Ponemah software (Data Sciences International) and analyzed to obtain beat-to-beat systolic, diastolic, and mean arterial pressures, which were averaged over consecutive 1-min intervals. The lights were on from 7:00 AM to 7:00 PM and off from 7:00 PM to 7:00 AM. Data collected within 1 h before and after the lights-on and lights-off transitions were excluded from the analysis.

### Myocardial infarction (MI) procedure

Myocardial infarction (MI) was performed by the Rodent Cardiovascular Phenotyping Core (RRID: SCR_022419) at the University of Pennsylvania. Mice were administrated analgesics (Meloxicam extended-release, 6 mg/kg, and Ethiqa-XR buprenorphine extended-release, 3.25 mg/kg) pre-operatively and 48 hours post-MI (Ethiqa-XR only, 3.25 mg/kg). The mice were anesthetized with ketamine (100 mg/kg) and xylazine (10 mg/kg) intraperitoneally. The mice were intubated and ventilated with a Harvard ventilator with a tidal volume of 200–400 µL. A left thoracotomy was performed through a small incision in the intercostal space between third and fourth ribs. MI was induced by permanent ligation of the left anterior descending coronary artery with an 8-0 silk suture.

### Echocardiography

Transthoracic echocardiography was performed using a Vevo 2100 Ultrasound System (Fujifilm VisualSonics Inc, Toronto, Ontario, Canada). Mice were lightly anesthetized by intraperitoneal administration of Avertin (2,2,2-tribromoethanol; Sigma-Aldrich, St. Louis, MO) at 0.005 mL/g body weight. After removal of hair from the anterior chest with a chemical depilatory, mice were positioned in the left lateral decubitus position on a warming pad to maintain a body temperature of 37°C. Ultrasound gel was applied to the chest, and adequate transducer contact was maintained without exerting excessive pressure. Two-dimensional B-mode images were acquired by manually positioning the ultrasound transducer to obtain parasternal long and short-axis images for assessment of left ventricular systolic function and regional wall-motion abnormalities following MI. Following imaging, mice were allowed to recover from anesthesia before being returned to their cages.

### Molecular Docking Simulation

The hydrated docking simulations were performed in Python using Autodock Vina^20,21^. AlphaFold PDB file containing the receptor coordinates for Human AT1R (AF-P30556-F1) was first modified with the reduce script^22^ ($ reduce -BUILD receptor.pdb), adding all missing hydrogen atoms. PDBQT file and grid parameters of the receptor were then generated using the python package Meeko ($ mk_prepare_receptor.py)^23^. Affinity maps and water map were finally computed with autogrid4 and the python Vina package mapwater.py, respectively. Ligands coordinates were obtained through PubChem (sdf files, 3D structures) and hydrogen atoms were first added with the python package Molscrub ($ scrub.py -I ligand.sdf -0 ligandH.sdf). The meeko script ($ mk_prepare_ligand.py -i ligandH.sdf -o ligand.pdbqt -w) was then used to add explicit water molecules to the ligands and generate the PDBQT files. The Docking affinities were calculated using Autodock Vina with Autodock 4 forcefield and the exhaustiveness set at 32, following Vina hydrated docking protocol (https://autodock-vina.readthedocs.io/). The computed docking was visualized in the software PyMOL (Schrodringer). Molecular interactions between AT1R, BT2, PF07328948 and Losartan were identified using the web tool PLIP (https://plip-tool.biotec.tu-dresden.de/plip-web/plip/index)^24^.

Molecular docking simulation in the inactive conformation of the receptor was performed using the Glide module within Schrödinger Maestro, and each ligand was first energy-minimized and ionization-corrected for physiological pH using the LigPrep tool. The orthosteric binding site was mapped using receptor grids centered on the co-crystallized ligands of the established inactive conformation AT1R crystal structure, PDB ID 4YAY, bound to the selective antagonist ZD7155, resolved at 2.9-Å resolution by X-ray crystallography. Because docking scores can be unreliable when comparing poses across structurally distinct receptor states, we focused our interpretation on the nature and quality of the molecular contacts each compound formed within the pocket, specifically hydrogen bonds, salt bridges, π-cation interactions, and hydrophobic π-π stacking, together with the overall shape and electrostatic complementarity between the ligand and binding site.

### AT1R BT2 Displacement Assay

Crude membranes were prepared from transiently transfected tetracycline-inducible Expi293F cells that had been incubated with a pcDNA-Zeotet0-AT1R construct as previously described^25^. Membranes from cells not transduced with the AT1R construct were prepared in parallel and used as controls. AT1R containing and control membranes were incubated with 1uM BT2 in 350ul of assay buffer containing 50mM Tris pH 7.4, 150 mM NaCl, 12.5 mM MgCl_2_, and 1x LB protease inhibitor for 30 minutes at room temperature with gentle shaking. Membranes were then pelleted by centrifugation and washed in 50 mM Tris (pH 7.4) three times to remove unbound BT2. Serial dilutions of Losartan in assay buffer were then added to the membranes for 90 minutes with gentle shaking. Lastly, membranes were pelleted by centrifugation, and the supernatant was collected for quantification of displaced BT2 by Liquid Chromatography-Mass Spectrometry (LC-MS). For LC-MS, 0.2 nmol of the standard F-BT2 was added to each sample to permit accurate quantification of BT2. Samples were then diluted in methanol and dried under a stream of nitrogen at 37^◦^C. Dried preparations were then resuspended in a 2:1 (v/v) methanol: water mixture and BT2 was quantified relative to the F-BT2 standard via LC-MS/MS using a Sciex QTRAP 6500^+^ mass spectrometer. Chromatographic separation was performed on an XBridge C18 analytical column (3.5 µm, 4.6 × 50 mm; Waters) connected to a guard column (VWR). The mobile phases were A, 98% H_2_O/2% acetonitrile (ACN) containing 0.1% formic acid, and B, 98% ACN/2% H_2_O containing 0.1% formic acid. The gradient was initiated at 0% B and maintained for 2 min, followed by an increase to 90% B over 10 min and then to 95% B over 1 min. The column was maintained at 95% B for 4.5 min for column washing, followed by a return to 0% B over 0.5 min and re-equilibration at the initial conditions for 7 min. The flow rate was 0.4 mL/min. The column oven and autosampler temperatures were maintained at 40°C and 4°C, respectively.

The optimized mass spectrometric parameters were as follows: ion spray voltage, 5500 V; entrance potential (EP), 10 V; collision cell entrance potential (CEP), 14 V; curtain gas, 35 psi; collision-activated dissociation (CAD), 10; source temperature, 650°C; and gas 1 and gas 2, 62 psi. Multiple-reaction monitoring (MRM) transitions were m/z 246.8 → 167.9 for BT2 and m/z 230.9 → 151.9 for F-BT2. Displacement curves were plotted in Graph Pad Prism 11. The presence of membrane AT1R was validated by western blot of the membrane fraction using the DYKDDDDK tag Mouse Mc antibody from Proteintech (cat:66008-4-Ig).

### Cell Culture and Transfection

Human embryonic kidney 293T cells were maintained in minimum essential medium (MEM) supplemented with 10% fetal bovine serum (FBS) and 1% penicillin/streptomycin (P/S) at 37°C and 5% CO₂. Transient transfections were performed using polyethylenimine (PEI) at a PEI:DNA ratio of 3:1.

### NanoBiT β-arrestin Recruitment Assays

HEK293T cells seeded in 6-well plates were transiently transfected with FLAG-AT1R, SmBiT-β-arrestin 1, and LgBiT-CAAX. Twenty-four hours after transfection, cells were plated onto clear-bottom, white-walled 96-well plates (Costar) at 100,000 cells/well in clear MEM supplemented with 2% FBS, 1% P/S, 10 mM HEPES, 1× GlutaMAX, and 1× Antibiotic-Antimycotic (anti-anti) (Gibco). The following day, media were aspirated, and cells were incubated with 60 μL of BT2 or vehicle for 25 min, followed by addition of 20 μL coelenterazine h diluted in Hanks’ balanced salt solution (HBSS; Gibco) supplemented with 20 mM HEPES to a final concentration of 2.5 μM for 5 min. Luminescence at 480 nm was measured at 37°C using a Berthold Mithras LB 940 plate reader. Three baseline reads were acquired before stimulation, followed by addition of 20 μL of 5X AngII for dose response curves. Luminescence responses were calculated as the change in signal over baseline and normalized to vehicle-treated controls.

### BRET β-arrestin Recruitment and Receptor Internalization Assays

For internalization assay, HEK293T cells seeded in 6-well plates were transiently transfected with receptor donor constructs, either AT1R-Rluc8 or V2R-RlucII, and 2xFYVE-mVenus. For β-arrestin recruitment assay, cells were transfected with V2R-RlucII and β-arrestin1-YFP. Twenty-four hours after transfection, cells were plated onto clear-bottom, white-walled 96-well plates (Costar) at 100,000 cells/well in clear MEM supplemented with 2% FBS, 1% P/S, 10 mM HEPES, 1× GlutaMAX, and 1× anti-anti. The following day, media were aspirated, and cells were incubated with 60 μL of BT2 or vehicle for 25 min, followed by addition of 20 μL coelenterazine h diluted in HBSS supplemented with 20 mM HEPES to a final concentration of 2.5 μM for 5 min. Signals were measured at 37°C using a Berthold Mithras LB 940 plate reader with 480 nm and 530 nm emission filters to detect donor and acceptor emission, respectively. Three baseline reads were acquired before stimulation, followed by addition of 20 μL of 5X AngII for dose response curves. BRET ratios were calculated by dividing the acceptor emission signal by the donor emission signal. Net BRET was calculated by subtracting the BRET ratio of vehicle-treated cells from that of AngII-treated cells.

### Calcium Influx Assay

HEK293T cells were transfected with FLAG-AT1R and, 24 h later, plated onto clear-bottom, black-walled 96-well plates (Costar) at 100,000 cells/well in clear MEM supplemented with 2% FBS, 1% P/S, 10 mM HEPES, 1× GlutaMAX, and 1× anti-anti. The following day, media were aspirated, and cells were incubated with 80 μL of 5 μM Cal-520 AM dye (AAT Bioquest) diluted in HBSS containing 20 mM HEPES for 60 min at 37°C. Cells were then treated with 20 μL of BT2 for an additional 30 min. The dye-containing solution was removed, and 80 μL of assay buffer was added to each well. Fluorescence was measured at 37°C using a BioTek Synergy Neo2 plate reader with excitation/emission wavelengths of 480/530 nm. Five baseline reads were acquired before stimulation, followed by addition of 20 μL of AngII. Calcium responses were quantified as F/F₀, where F represents the fluorescence signal following AngII stimulation and F₀ represents the fluorescence signal with vehicle addition.

### Statistical analysis

Statistical analyses were performed using GraphPad Prism version 11. Data are presented as mean ± SD. Comparisons between two groups were performed using a two-tailed unpaired Student’s t test unless otherwise specified. Comparisons among multiple groups were performed using one-way or two-way ANOVA followed by the multiple-comparisons test specified in the corresponding figure legend. The number of mice represents the biological sample size (n). A P value < 0.05 was considered statistically significant.

---

**Supplemental Figure 1.**
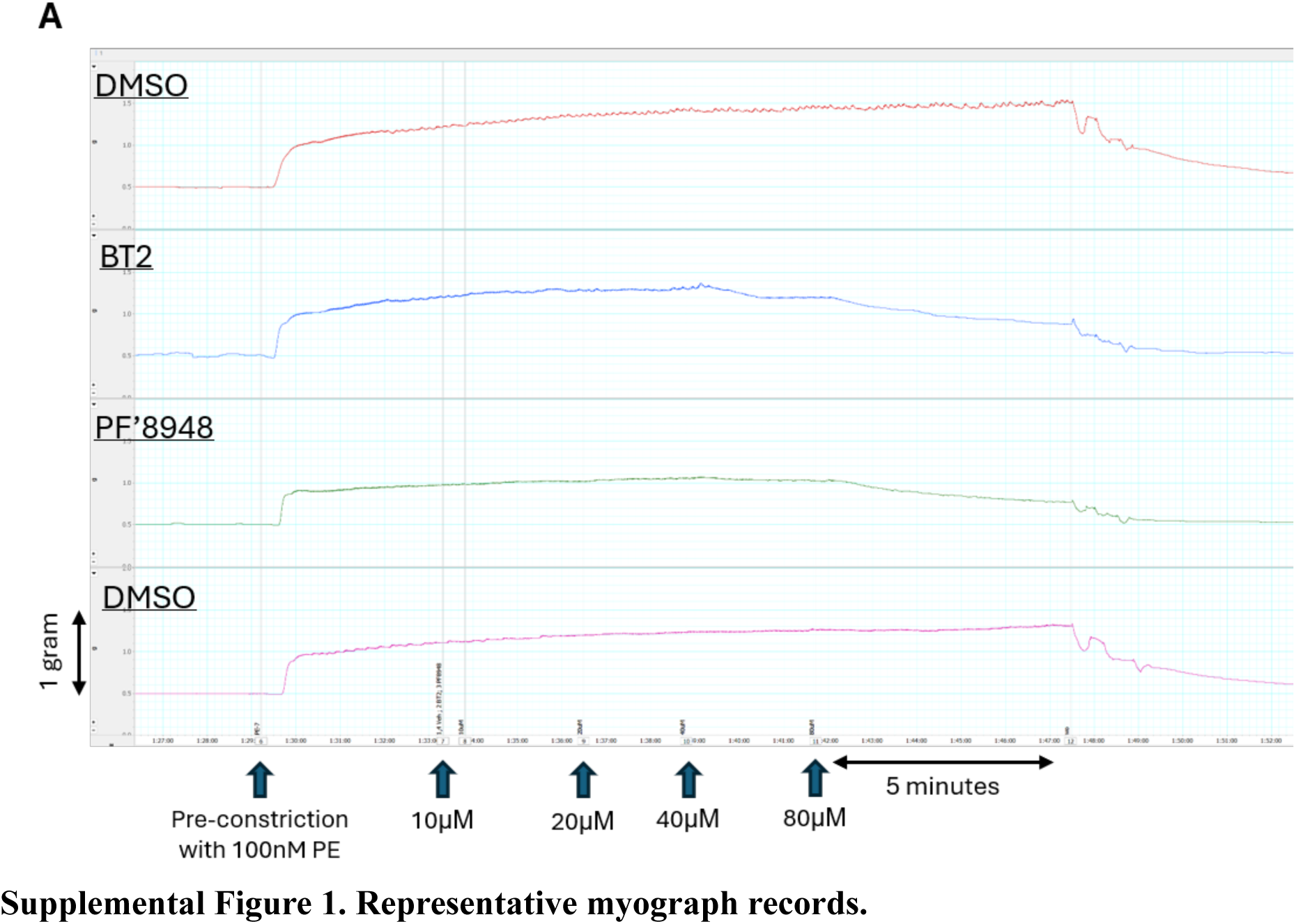
Representative myograph records.

**Supplemental Figure 2.**
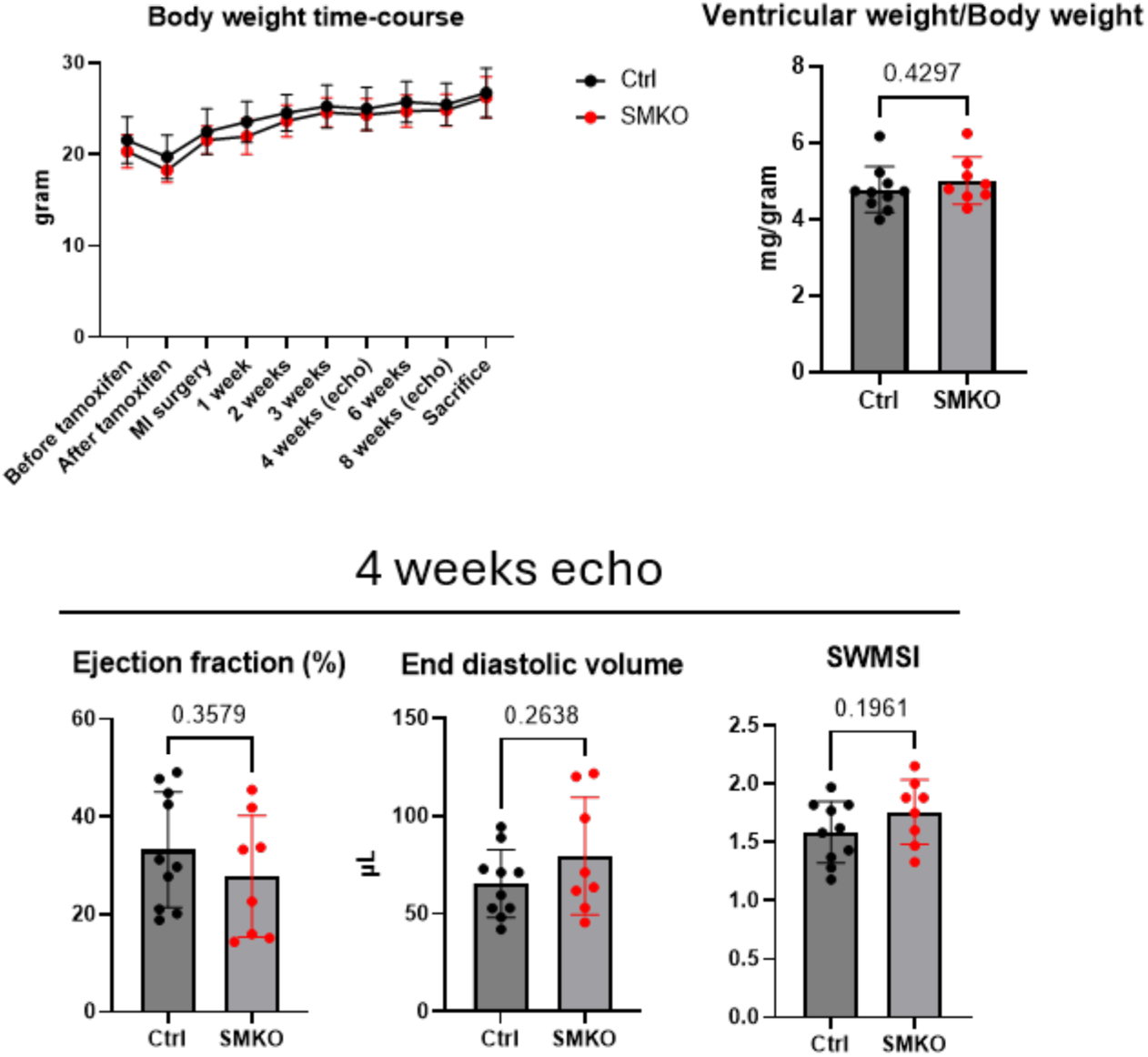
Body weight, heart weight, and echo data 4 weeks after myocardial infarction. Unpaired t-test with Welch’s correction was performed to compare SMKO and Ctrl.

**Supplemental Figure 3.**
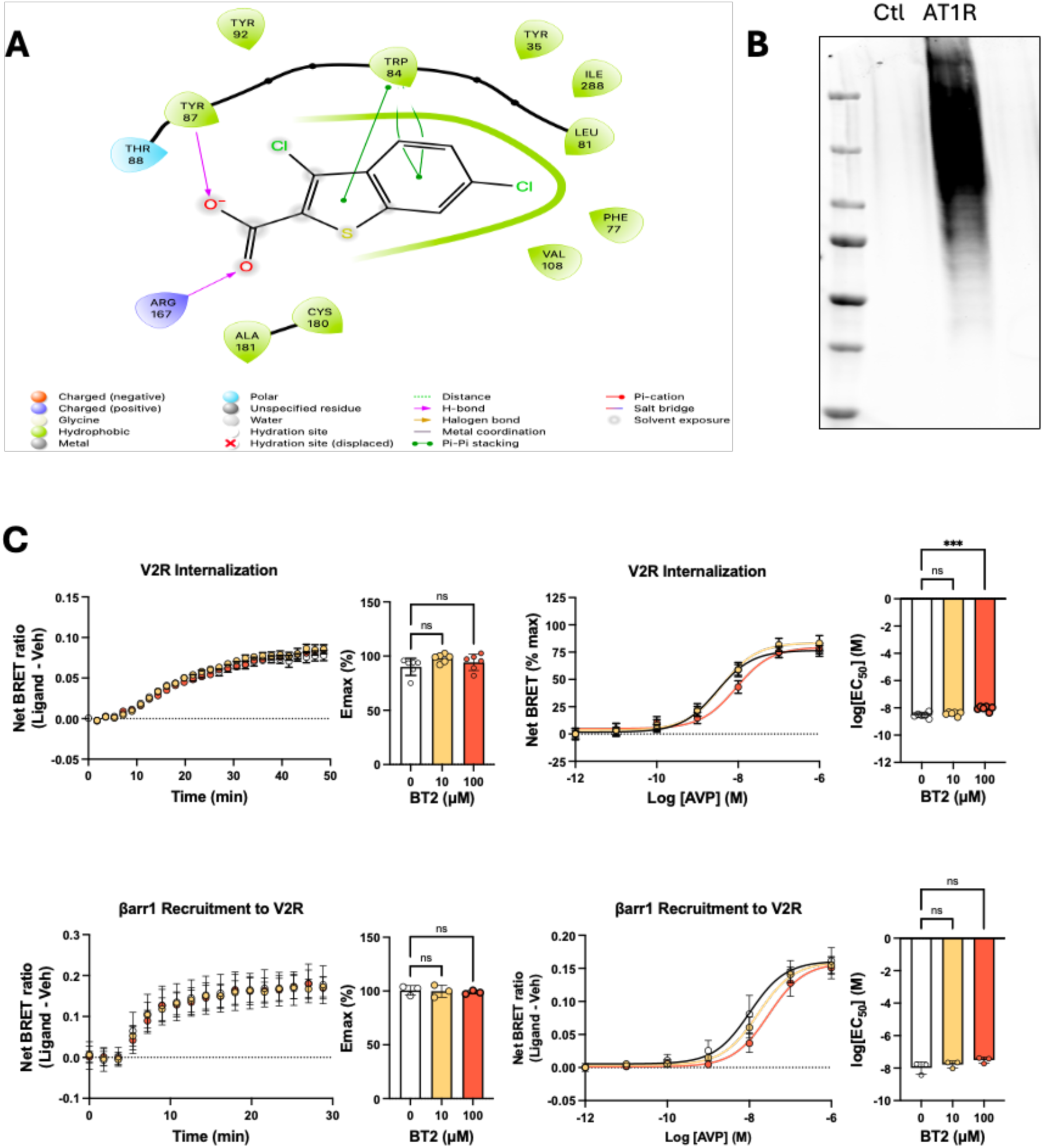
**A)** Predicted docking of BT2 in the orthosteric pocket of AT1R in the inactive conformation. **B)** Western blot for AT1R of control (Ctl) Expi293 membranes and those overexpressing AT1R (AT1R). **C**) HEK293T cells expressing V2R were pretreated with BT2 or vehicle before stimulation with arginine vasopressin (AVP). V2R internalization was assessed by BRET between V2R-RlucII and the endosomal marker 2×FYVE-mVenus (n = 6 independent experiments), and β-arrestin 1 recruitment was assessed by BRET between V2R-RlucII and β-arrestin 1-YFP (n = 3 independent experiments). Data are mean ± SEM. Statistical comparisons were performed by one-way ANOVA with Dunnett’s post hoc test comparing BT2 versus vehicle-treated cells; ***p < 0.001; ns, not significant.

